# Strain maps characterize the symmetry of convergence and extension patterns during Zebrafish gastrulation

**DOI:** 10.1101/407940

**Authors:** Dipanjan Bhattacharya, Jun Zhong, Sahar Tavakoli, Alexandre Kabla, Paul Matsudaira

**Affiliations:** Center for BioImaging Sciences (CBIS), Department of Biological Sciences, National University of Singapore, Singapore 117543; MechanoBiology Institute, National University of Singapore, Singapore 117411; Singapore-Massachusetts Institute of Technology Alliance of Research and Technology (SMART), National University of Singapore, Singapore 117548; Department of Engineering, University of Cambridge, Cambridge CB2 1PZ, United Kingdom; Institute FIRC of Molecular Oncology (IFOM), Milan 20139, Italy; Harvard University, Department of Stem Cell and Regenerative Biology, Harvard University, Cambridge 02138, USA; Boston Children’s Hospital, Boston 02115, USA; Harvard Stem Cell Institute, Harvard University, Cambridge 02138, USA

**Keywords:** gastrulation, convergence and extension, collective migration, left-right symmetry, zebrafish, mechanics, tectonics, strain map, curl

## Abstract

During gastrulation of the zebrafish embryo, the cap of blastoderm cells organizes into the axial body plan of the embryo with left-right symmetry and head-tail, dorsal-ventral polarities. Our labs have been interested in the mechanics of early development and have investigated whether these large-scale cells movements can be described as tissue-level mechanical strain by a tectonics-based approach. The first step is to image the positions of all nuclei from mid-epiboy to early segmentation by digital sheet light microscopy (DSLM), organize the surface of the embryo into multi-cell spherical domains, construct velocity fields from the movements of these domains and extract 3D strain rate maps. Tensile/expansive and compressive strains in the axial and equatorial directions are detected during gastrulation as anterior and posterior expansion along the anterior-posterior axis and medial-lateral compression across the dorsal-ventral axis corresponding to convergence and extension. In later stages in development are represented by localized medial expansion at the onset of segmentation and anterior expansion at the onset of neurulation. Symmetric patterns of rotation are first detected in the animal hemispheres at mid-epiboly and then the vegetal hemispheres by the end of gastrulation. By analysing the temporal sequence of large scale movements, deformations across the embryo can be attributed to a combination of epiboly and dorsal convergence-extension.

**Significance:** Strain is an emergent property of tissues that originates from the mechanical coupling of cell-cell and cell-substrate interactions, individual cell shape changes, and cell level forces. By imaging the positions of nuclei from mid-epiboly to early segmentation of the zebrafish embryo we are able to calculate three types of strain maps by a plate tectonics based method. The regions of expansive and compressive axial and equatorial strain correspond to areas undergoing convergence and extension, a major step in the formation of the embryonic body plan as well as the formation of somite and head structures. The most striking signatures of strain are: 1. the bilateral symmetry of linear strain across the anterior-posterior, dorsal-ventral axis during gastrulation, 2. the complementary counter-rotational strains or curl in the animal hemisphere at mid epiboly, and 3. a divergence or saddle point in the region of the dorsal organizer, head-trunk boundary. These strains represent a general method to describe large-scale tissue-level mechanics not only of embryonic development but also tissue homeostasis and disease.

## Introduction

A hallmark of zebrafish early development is gastrulation when the hemispherical shield of the blastoderm rearranges by the collective cell migrations and rearrangements of convergence and extension into the axial and symmetric body plan of the embryo. These large-scale cell movements and patterns are coordinated biochemically by a chemical gradient of morphogens such as BMP, which regulates convergence and extension at the cell-to-cell level via a calcium-dependent cell adhesion pathway (1). Together with Nodal, these two morphogens are the minimum requirement for establishing the body axis. (2) Thus, morphogenesis is the product of cell-to-cell signalling at a local scale regulated by gradients of morphogens at the whole embryo scale.

Chemical signals also coordinate changes in cell shape, cell-cell-matrix adhesions, and cell arrangements (3) of morphogenesis. These cell-level changes result from well-studied forces including molecular motors and cytoskeletal dynamics at cell adhesions (3–7). When these forces are coordinated across cells, tissue-level strains and tension are generated which contribute to the dynamics of embryonic tissue folding and dorsal closure as well as wound healing and other tissue-scale movements (8).

While the biochemistry, cell biology, and cell-level mechanics have explained key features of morphogenesis, the role of stress and strain in morphogenesis is now being explored. Recent advances in imaging and image processing approaches can now extract from whole embryo time-lapse images several key signatures of large-scale dynamics such as velocity fields, cell density changes, and 2D tissue deformations (9,10). However, to detect and describe the mechanics of tissue-derived forces in the context of an entire embryo requires large-scale 3D *in toto* imaging with: 1. high speed to capture developmental dynamics, 2. high resolution to resolve individual cells, and 3. high penetration depth to image whole organisms. While confocal and multi-photon microscopy have contributed fundamental information on early development, SPIM/DSLM based microscopes have emerged as the instrument of choice in studies of Zebrafish or Drosophila embryogenesis (10, 11). With the temporal and spatial resolution and reduced photo-bleaching of SPIM/DSLM, it is possible to track every cell during early development (10). As a result, the technical imaging challenges are replaced by the computational challenge of extracting the tissue-level mechanics from very large image datasets.

To detect and characterize the large-scale tissue dynamics during gastrulation of the zebrafish embryo, we have adapted the 2D tectonics-based analysis of tissue-level strain to the 3D analysis of deformation over the whole embryo. The resulting strain rate maps reveal the time-course and location of areas undergoing compaction or expansion, i.e. linear strain, during well-known morphogenetic movements including convergence and extension, neural plate formation, and somite formation. In addition to axial and equatorials strains, new features such as rotational strain or curl appear during convergence and extension as well as the left-right laterally symmetric presence of strain across the embryonic axis. These strains confirm that the cell-level dynamics and interactions of epiboly and gastrulation are mechanically coupled at the tissue-level and conform to the bilateral symmetry established in the embryo after fertilization.

## Results

To detect and characterize the mechanical strain during gastrulation, first we imaged all cells continuously from 75% epiboly (8 hpf) to 3 somite stage (11 hpf) (Figure 1a, Movie 1), calculated their velocity fields, and color-coded the direction of the velocity fields (Figure 1). The velocity fields cover the entire surface of the embryo and capture the major morphogenetic movements in three principal directions: 1. epiboly as ventral translocation (red), 2. convergence as lateral translocation (blue), and 3. extension as anterior translocation (green). These movements persist through developmental stages 90% epiboly (9 hpf), 100% epiboly (10 hpf) and 1-somite (10.5 hpf) (Fig 1B, 1C). In the ventral view of the 10 hpf bud stage embryo, the extending tail bud region is seen extending anteriorly from the vegetal pole.

**Figure 1.**
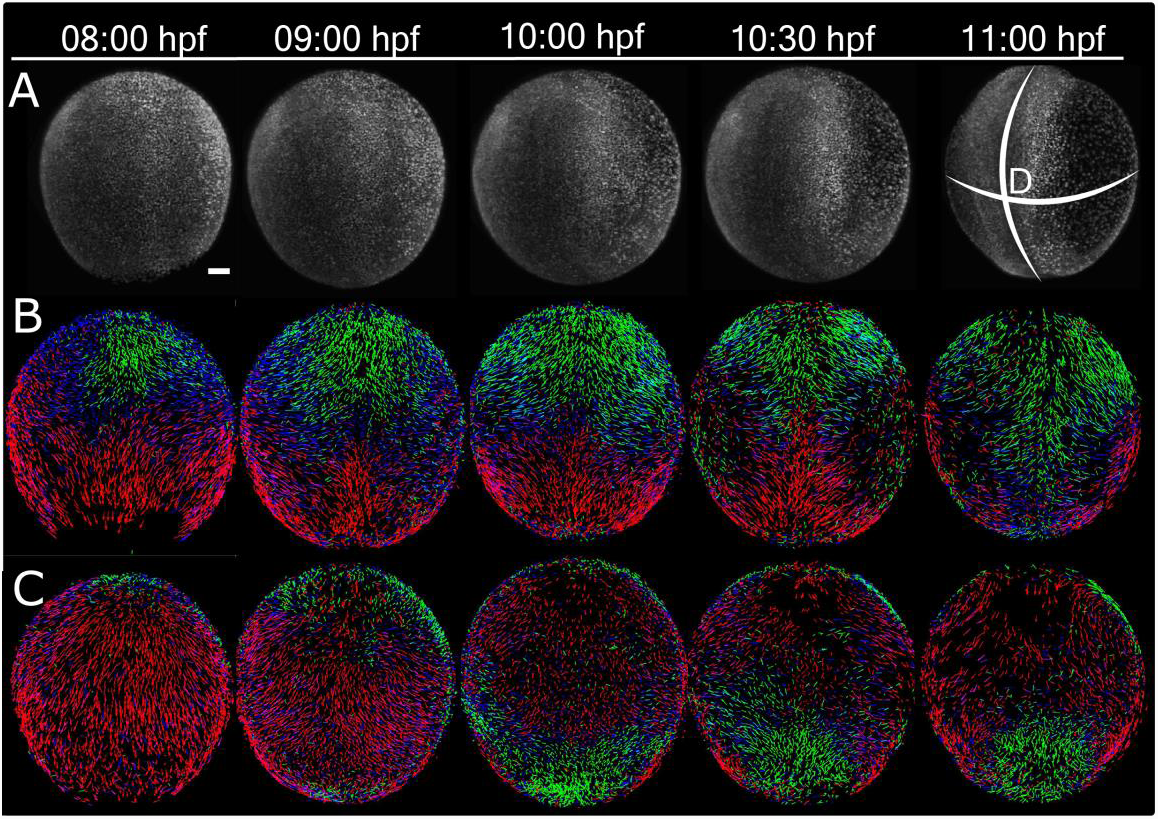
Velocity field maps of zebrafish from mid-gastrula to early segmentation stages. (A) Dorsal views of GFP-histone H2B-tagged embryos at 8 hpf (75% epiboly), 9 hpf (90% epiboly), 10h pf (100% epiboly), 10.5 hpf (1 somite), 11 hpf (1-3 somite stages) respectively. Velocity field maps of the dorsal (B) and ventral (C) views of the velocity field maps. The length of the velocity vector represents the magnitude of the domain velocity over a 2-minute interval. The color of the vectors represents the directions: Green towards animal pole, Red towards Vegetal pole, and Blue color represents the medio-lateral convergence or divergence. For all views, the embryo is oriented with anterior (A) and posterior (P) to the top and bottom respectively. Scale bar-100μm.

To identify the location and magnitude of tissue deformation over the embryo, we then calculated 3D tissue deformation or strain by determining the 3D strain rate tensor in three principal directions: the anterior-posterior (AP), media-lateral (ML), and radial (r) directions (supplementary materials). In our analysis, we display only the AP (axial) and ML (equatorial) as these directions align with the major changes during morphogenesis (Figure 2B). Several patterns of strain are present in the maps of the dorsal and ventral hemispheres of the embryo. First from 8-10 hpf, the dorsal hemisphere is undergoing M-L compaction while simultaneous A-P expansion is restricted to an equatorial band around the dorsal but not ventral half of the embryo. Second, the dorsal M-L compaction is accompanied by extensive M-L expansion in the ventral hemisphere. Third, by 10:30 hpf the dorsal compaction narrows to the A-P body axis but is now flanked by a prominent a pair of M-L expansions (asterisk). This expansion is transient, lasting until 11 hpf and diminishes afterwards (n = 8). At this time the ventral hemisphere compacts along the A-P axis as the body axis extends into the ventral side to form future head and tail of the embryo.

**Figure 2.**
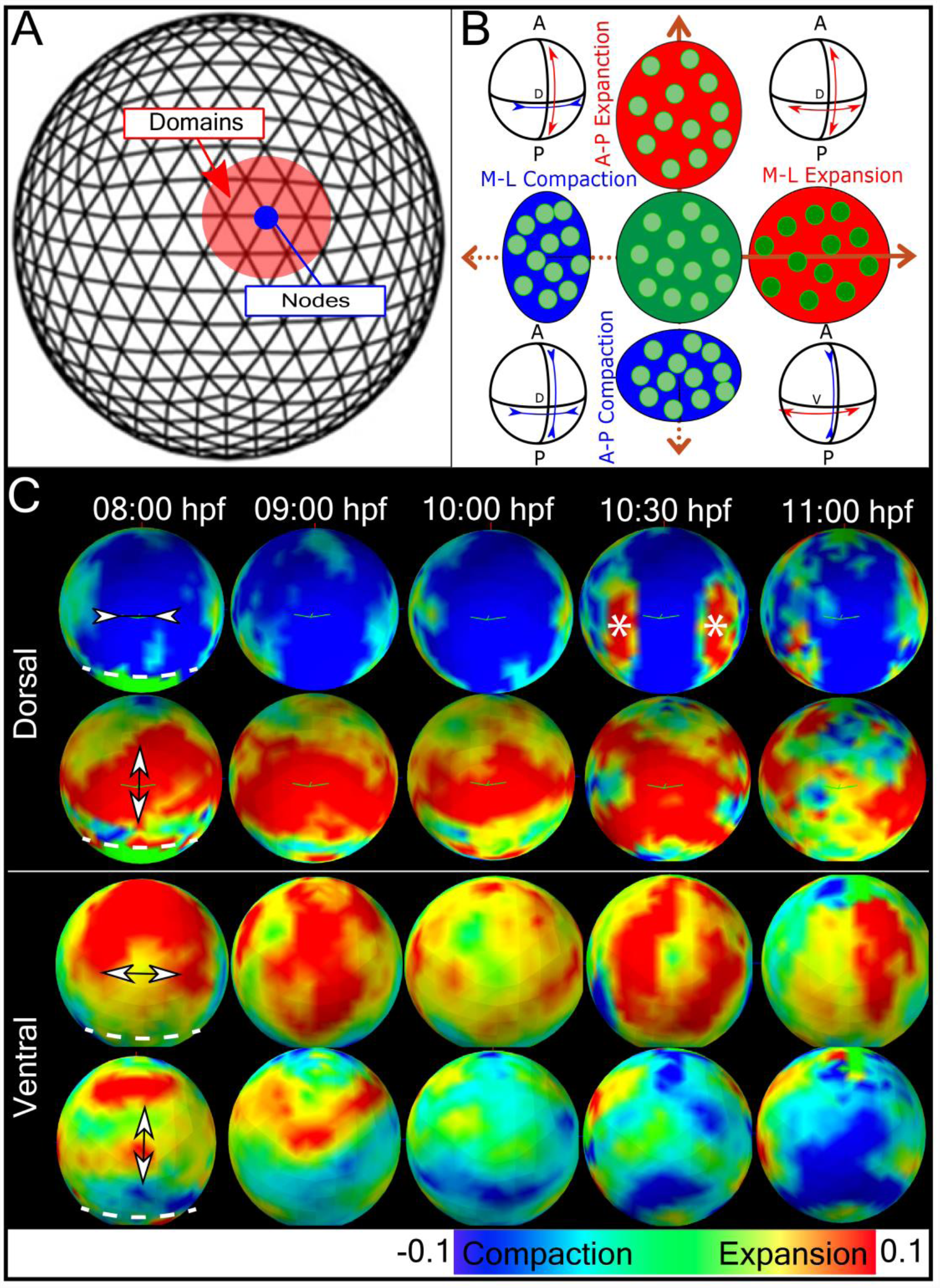
Whole embryo strain rate maps during morphogenesis. (A) Diagram of the spherical embryonic surface discretized as a mesh defined by nodes and domains for computing the strain-map. (B)The orthogonal components of strain along A-P and M-L directions. Expansion (red) and compaction (blue) of domains are shown oriented along A-P and M-L axis. For simplicity, strain in radial direction was not shown here. The strain here and the rest of the study are obtained from strain rate integrated over a 30-minute interval centred at the indicated time point. (C) Experimentally derived strain-maps along Medio-lateral (M-L) and Anterior-posterior (A-P) directions. Dorsal and ventral views of strain maps along M-L and A-P directions at five developmental stages. The embryo is oriented with anterior and posterior at the top and bottom respectively. Expansion state in both side of the dorsal line at 10:30 hpf, is marked with “*” symbol.

To understand the time course in the evolution of strain, we constructed kymographs of the three strain components in two important regions of the body: the circumference around the equatorial plane (Figure 3B,C,D) and the axial circumference along the AP axis (Figure 4B,C,D). The kymograph of M-L-directed strain shows a persistent dorsal compaction and ventral expansion (Figure 3B) and is accompanied by the prominent dorsal expansion in the AP direction (Figure 3C). The M-L and A-P strains are bilaterally symmetric across the 0° dorsal point. Finally, radial strain (Figure 3D) is initially detected as compaction in areas midway between the dorsal and ventral sides of the embryo but by 10:30 hpf the dorsal point of the embryo shows prominent expansion (Figure-3D).

**Figure 3.**
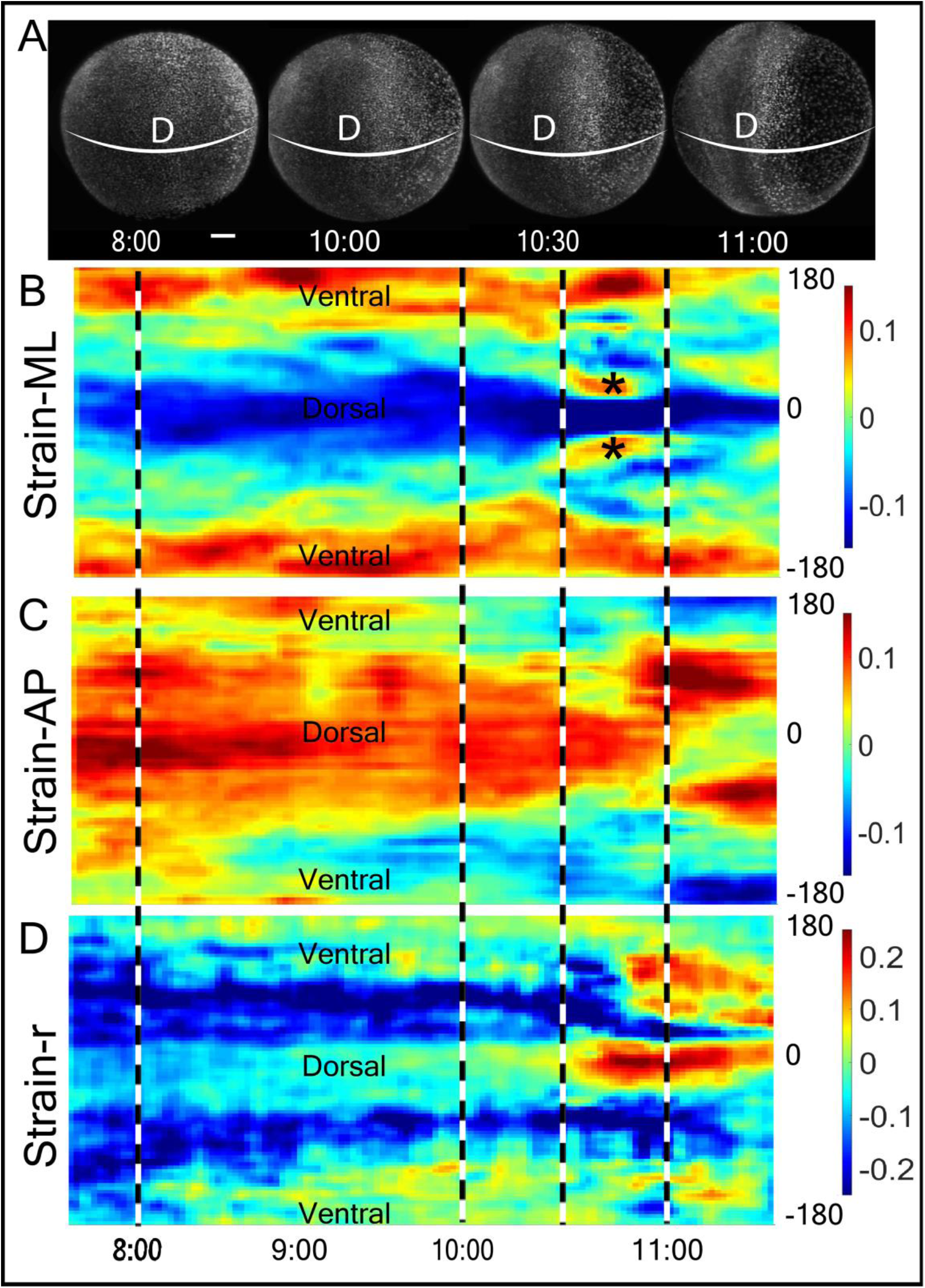
Time evolution of strain around the equatorial plane along M-L, A-P and radial (‘r’) directions. (A) The dorsal image of the embryo at 8, 10, 10.5 and 11 hpf of zebrafish development. Strain is calculated along the embryo equator (white band). Scale bar-100μm. (B,C,D) Kymograms of strain along (B) M-L, (C) A-P, and (D) radial, ‘r’, directions and from 7.5 to 11.5 hpf (x-axis). The Y-axis measured strains around the circumference (white band in (A) of the embryo) from −180° (Ventral side) to 0° (Dorsal side) to 180° (ventral side) in the anticlockwise directions. Dashed lines correspond to stages in (A). Expansion state in both side of the dorsal line at 10:30 hpf, is marked with “*” symbol.

**Figure 4.**
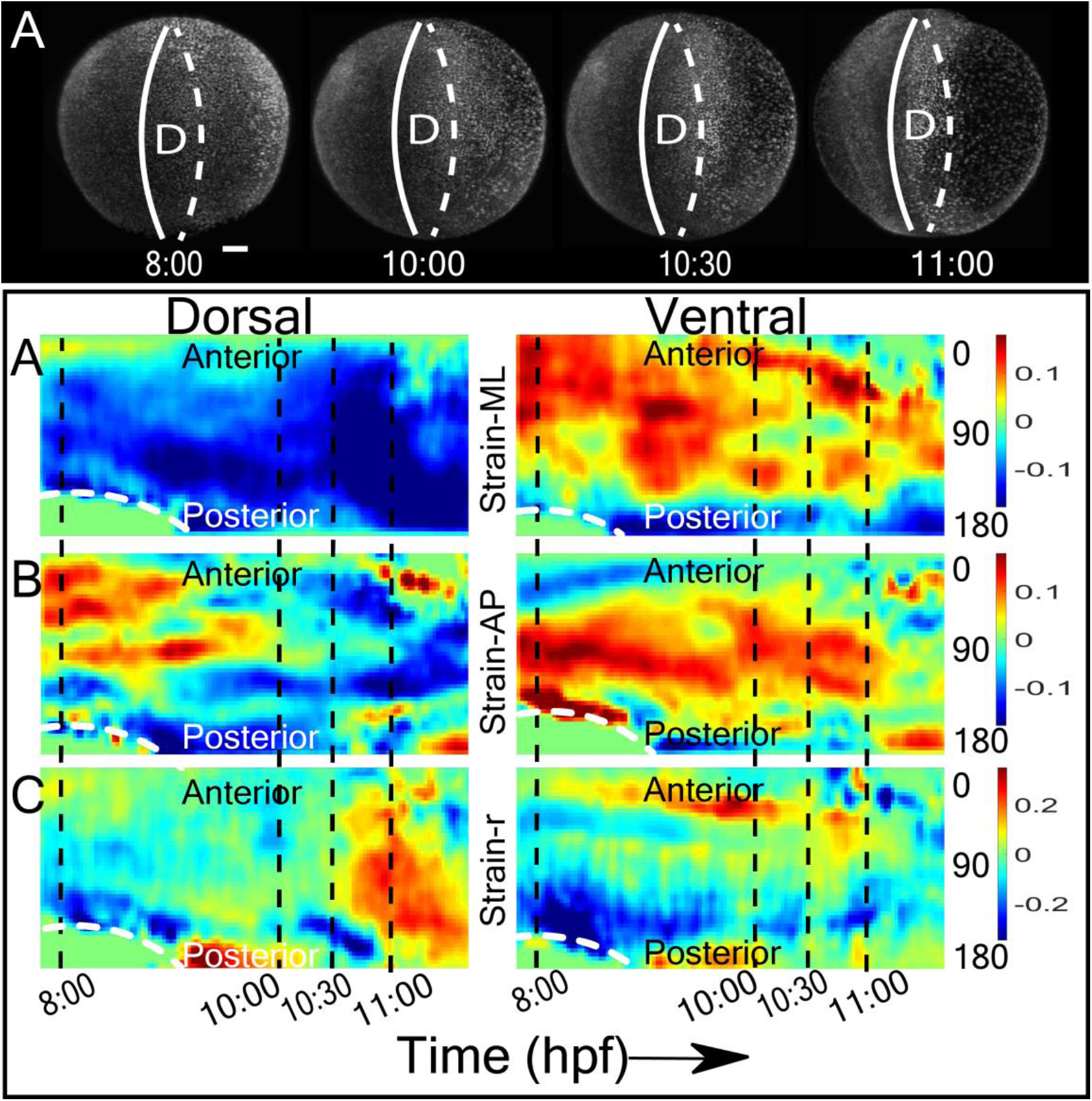
Time evolution of strain around the axial plane in the M-L, A-P and radial (‘r’) directions. (A) The dorsal view of the embryo at 8, 10, 10.5 and 11 hpf of zebrafish development. Strain is calculated along the embryo anterior-posterior axis corresponding to the dorsal (solid white band) and ventral (dashed white band) midline of the embryo. Scale bar-100 µm. (B,C,D) Kymograms of dorsal (left panels) and ventral (right panels) strain in the M-L (B), A-P (C), and radial, ‘r’ (D) directions and from 7.5-11.5 hpf (x-axis). The Y-axis measures strain around the anterior-posterior plane (white band in (A) of the embryo) from the anterior (0°) to the posterior side (180°). Dashed lines correspond to stages in (A). The white dotted line across the posterior side between 8-9 hpf indicates the margin of dorsal closure.

We then analysed strain along the body axis of the embryo (Figure-4). Figure-4(A) indicates the continuous compaction along M-L direction at dorsal site. We observe that the dorsal compaction becomes stronger, extending from the equator plane towards 2 polar regions. Again, the kymograph shows this M-L compaction comes along with A-P expansion during epiboly, followed (10.5+ hpf) by an increase in radial strain at the same spatial position. This site also coincides with the place of early somite as well as the neural tube.

A deformation or change of the domain, i.e strain trace, can be calculated from the divergence of the strain rate tensor over a time period (set at 30 minutes). The resulting strain trace map shows large-scale change in the ventral hemisphere that extends medially and dorsally prior to 100% epiboly, followed by change along the dorsal body axis. These tissue deformations are inhomogeneous over the embryo and during morphogenesis.

Having identified the patterns of strain along the main embryo axes, we then visualized the amount of rotation in the velocity field (magnitude of the anti-symmetric part of the velocity gradient tensor), Figure-6. The resulting curl maps at 60%, 75%, 100% epiboly, bud, and first somite stages indicate distinct rotational strain of the embryonic tissues in positive, clock-wise (red) or negative, anticlockwise (blue) directions and cover regions of the embryo that lie within the left/right, anterior/posterior quadrants of the embryo. As early as 60% epiboly, the maps show counter-rotational strain in the left and right sides of the anterior pole. When viewed from the dorsal or ventral poles, the left-side strain rotates counter-clockwise while the right-side strain rotates clockwise. Both curl strains correspond the anterior left and right quadrants of the embryo and extends between the dorsal and ventral poles. Anterior left-right strain is detected until 1st somite stage when the regions of left-right strain move midway between the anterior and posterior poles. The posterior half of the embryo shows zero curl (green) except of lip of positive curl near the posterior pole on the dorsal surface. By the completion of epiboly, counter-rotational strains have appeared in the posterior half of the embryo and this strain pattern persists to segmentation. In contrast to curl strain in the left-right halves of the embryo, the curl strain is zero at the dorsal and ventral poles.

## Discussion

In this study we have detected and quantified three types of tissue deformation during epiboly and gastrulation: anterior-posterior (AP) and the medio-lateral (ML) strain, radial strain, and curl. Figure 2 presents a schematic representation of the deformation patterns. Notable features of these deformations are their long-range coupling and symmetry. An example of long range coupling is first seen during epiboly (7-10 hpf) when the cells migrate in all directions to cover the yolk. This expansion is visible from the anterior view of the embryo as A-P strain (Fig 2). However, before the cells reach the vegetal pole the strain maps also detect signatures of strain corresponding to convergence as ML compaction over the entire dorsal hemisphere. This compaction is accompanied by a broad equatorial band of expansion in the AP direction centered at the midpoint of the dorsal axis. The dorsal midpoint corresponds to the boundary between head and trunk structures. Interestingly, these lateral displacements of tissues on the dorsal side are accompanied on the ventral side by an opposite strain pattern (Fig 4). Cells converging along the dorsal side must be balanced by cells expanding somewhere else, as observed on the ventral side during and after epiboly. Also, tissues tend to expand in AP on the dorsal side (although this effect is diminished after the end of epiboly), and converge on the ventral side. Thus, the major morphogenetic movements of epiboly and convergence/extension appear to be mechanically coupled over the dorsal and ventral surfaces of the embryo.

In addition to the expansion and compression of these movements is the bilateral left/right symmetry that is evident in the ML strain kymographs (Fig 3B,C) during epiboly and continuing past gastrulation. This symmetry is defined by the animal/vegetal and dorsal/ventral axis from 8-10.5 hpf. Perhaps the most striking example of the symmetry in the strain maps is seen at the first appearance of a somite (10.5 hpf). There is a dorsal band of ML compaction which is flanked by a short-lived (10.5-11 hpf) and narrow region of ML expansion (asterisk). This “hot spot” region spatially coincides with the paraxial mesoderm, also known as presomitic or somitic mesoderm, which commits to the formation of somites (16). However, this local strain precedes the time when morphogenesis corresponding to somatogenesis occurs and suggests that paraxial mesoderm expansion is a possible biomechanical signal for somatogenesis before apparent morphology formation. The correspondence between events on opposite side of the embryo illustrates that strain is bilaterally symmetrical and equally generated on the left and right sides of the embryo.

In contrast to linear strain that is expressed along the surface of the embryo, we are able to detect radial strain during morphogenesis (Fig 3D, 4D). The most prominent deformations arise at 11 hpf when the neural structures are forming along the dorsal midline and in the ventral hemisphere. The ventral regions are undergoing significant expansion especially in areas corresponding to the head and tail structures. Thus, the radial strain corresponds to well characterized events in neurulation.

A third and particularly interesting type of deformation that we have characterized is rotational strain or curl. Curl appears in the dorsal hemisphere during epiboly and then in the ventral hemisphere when epiboly continues to the ventral pole. There are two striking aspects of the location and pattern of curl. First, the direction of curl is consistent with a large-scale integration of convergent movements toward the embryonic axis and extension along the embryonic axis in anterior and posterior directions into a unified rotational movement. Second, the direction of curl on the left and right sides of the embryo are opposite and complementary. When drawn on the surface of a sphere, these patterns of deformation clearly define the quadrants associated with the rotational patterns (Figure 7). These patterns reflect the bi-lateral symmetry of the cellular flows and coordination of morphogenetic movements over very large areas of the embryo.

Finally, a curious feature of convergence and extension that emerges from our analysis of the velocity fields is a divergent point or saddle at the dorsal midpoint from where cells translocate either anteriorly or posteriorly. In the curl maps, this divergence point corresponds to the region where the domains of positive and negative curl intersect. This point of divergence may have a functional significance. It corresponds to the site of ingression during epiboly and the boundary between head and trunk structures in the embryo. Our maps do not reveal the mechanism that directs cells to move or expand in opposite directions. We are investigating possible structural and molecular explanations for divergent cell migration.

In conclusion, the strain maps and 3D tissue tectonics are the first step to quantify the global biomechanics of developing embryos. To fully understand the force or tension within the tissue, the lack of validated methods to measure real time tissue stiffness remains a challenge. However, it has been reported the cell density is correlated with local tissue stiffness (20), and thus offers us an opportunity to gain an insight of relative tensions among tissues. A resulting tissue deformation-model can act as a mechanistic model to understand the coordinated activities of morphogen gradients, signalling pathways and physical forces that drive embryo morphogenesis.

## Materials and Methods

### H2B-GFP mRNA

H2B-GFP mRNA was transcribed (mMESSAGE mMACHINE^®^ SP6 Transcription Kit, Thermo Fisher Scientific) from a plasmid (a generous gift from Lora Tan) linearized with NotI restriction enzyme.

### Embryo Microinjection

Wild-type Zebrafish embryos were collected in the aquarium at 1-cell stage and immediately injected with 9.7 nL of H2B-GFP mRNA (150 ng/μL). Each embryo was kept at 28.5°C in the incubator. At 8 hours post-fertilization (hpf), the embryos were dechorionated, embedded in 0.4% low melting point agarose within a cleaned FEP tube, and then positioned in the DSLM for live imaging. The bottom of the FEP tube was sealed with 2% agarose solution.

### FEP Tube Cleaning

Prior to imaging, 1 mm inner diameter Fluorinated Ethylene Propylene (FEP) tubes (Bohlender GmbH) were cleaned by flushing with 1M NaOH, ultra-sonication for 10 min with 0.5 M NaOH solution, flushing with 70% EtOH and ultra-sonication for 10 mins in 70% ethanol solution. The cleaned tubes were flushed and stored in ddH2O.

### Live Embryo Imaging

Cleaned PEP tubes containing embryos injected with H2B-GFP mRNA were transferred into the imaging chamber of a custom-built Multi-View DSLM system (11). The chamber was held at 28.5 °C. The injected embryo was illuminated with a 488 nm laser and imaged with a Hamamatsu CMOS camera at 50 ms exposure/frame. A Multiview dataset consists of 450 mm stack with 2 mm spacing in four orthogonal directions (0°, 90°, 180°, 270°). 3D datasets were collected with two independent light sheets for excitation and single detectors in the Multiview mode, at 2 min intervals from 7.5 hpf to 12 hpf.

### Image Processing

Each experiment generates around 2 TB stack of TIFF image files which are processed with a MATLAB library (MathWorks, Inc) (10) to identify and track all nuclei in the embryo. The workflow includes: 1. 3D image reconstruction from 2D stacks, 2. image segmentation and cell identity registration using iterative thresh-holding method, 3. post-segmentation spatial filtering on intensity distribution, intensity value, anisotropy and connectivity, 4. temporal filtering that recovers false negatives and removes false negatives with reference to the nearby time points, and 5. 3D-cell tracking program adopting the nearest-neighbor algorithm.

### Strain maps

To calculate 3D strain we modified a previously described tectonics-based method for measuring strain in 2D (9). The general approach is to group all cells into overlapping spherical domains, calculate the velocity field gradients of the domains, and then extract the strain from the tensors of the velocity field gradient matrix. Because the early embryo is a sphere and to simplify the calculations, we start by mapping the embryo onto a unit sphere where the radius r0 is set to 1. Then we discretise the spherical embryo surface into 2000 equally spaced points or “nodes” (Figure 2A). Each node, 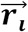, lies at the vertex of a triangular mesh and its location is described in polar coordinates 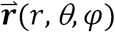where r is radial distance from the embryo centroid, θ is polar angle or elevation, and *φ* is azimuth. For each time point, t, we define local domains around each node by selecting all the cells within a certain distance from the node center. In this study, the radius of these domains is set to cover 110 microns which is small enough to capture relevant spatial patterns, while providing enough statistics to calculate the local strain rates. At 10 hpf, the domains would contain 10-250 cells. Once the domain is fixed, we obtain its local velocity field, **v,** from the mean of the cell velocities within a domain.

The velocity field is then analysed to extract the local strain rates, or the rate of tissue deformations (9). To calculate the strain rate, we differentiate the velocity field spatially, forming a 3×3 velocity gradient tensor 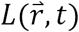:

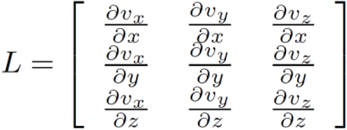

This tensor is then separated into its symmetric and anti-symmetric components. The symmetric part quantifies local deformations as the strain rate tensor:

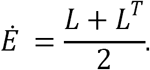

The tensor’s eigenvalue components are the magnitude of strain rates along the corresponding eigenvectors (r,θ,ϕ), and the sum of the diagonal components or trace of this matrix describes the total volume change of the domain. Because the major changes in development occur along the embryonic axis (A/P, Dorsal/Ventral) and equator (left/right), we decomposed all the vectors into anterior-posterior (A-P), medio-lateral (M-L), and radial (R) directions. The accumulated deformation or strain rates over time give rise to the magnitude of strains, in our case, we chose the time window of 30 minutes to filter random noise and highlight significant signals. The strains are colour coded where expansion is red, indicating diverging velocity fields or an overall increase in inter-nuclei distances while blue represents converging velocity fields or the cell cluster in the domain compacts and gets denser (Figure 2B).

In addition to linear strain, the anti-symmetric part of the tensor is the spin matrix and contains information about the local rotation or curl of the domain (Figure 6):

**Figure 5:**
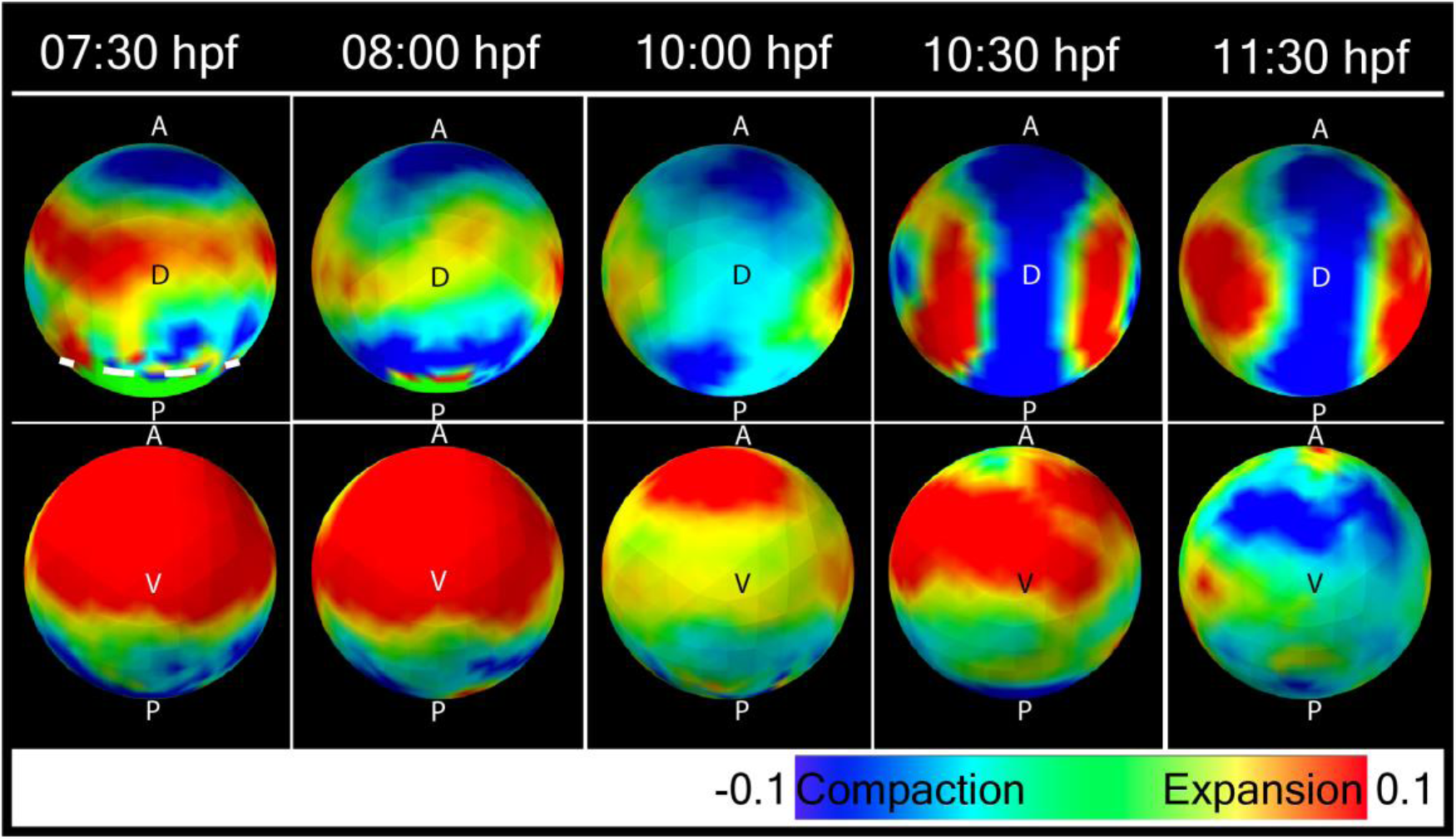
Divergence of velocity field (strain trace) at 7:30 (60% epiboly), 9 (90% epiboly), 10 (bud, 100% epiboly), 10.5 hpf (1-somite) and at 11:30 hpf. The white dotted line in the posterior side before 100% epiboly, indicates the margin of dorsal closure.

**Figure 6.**
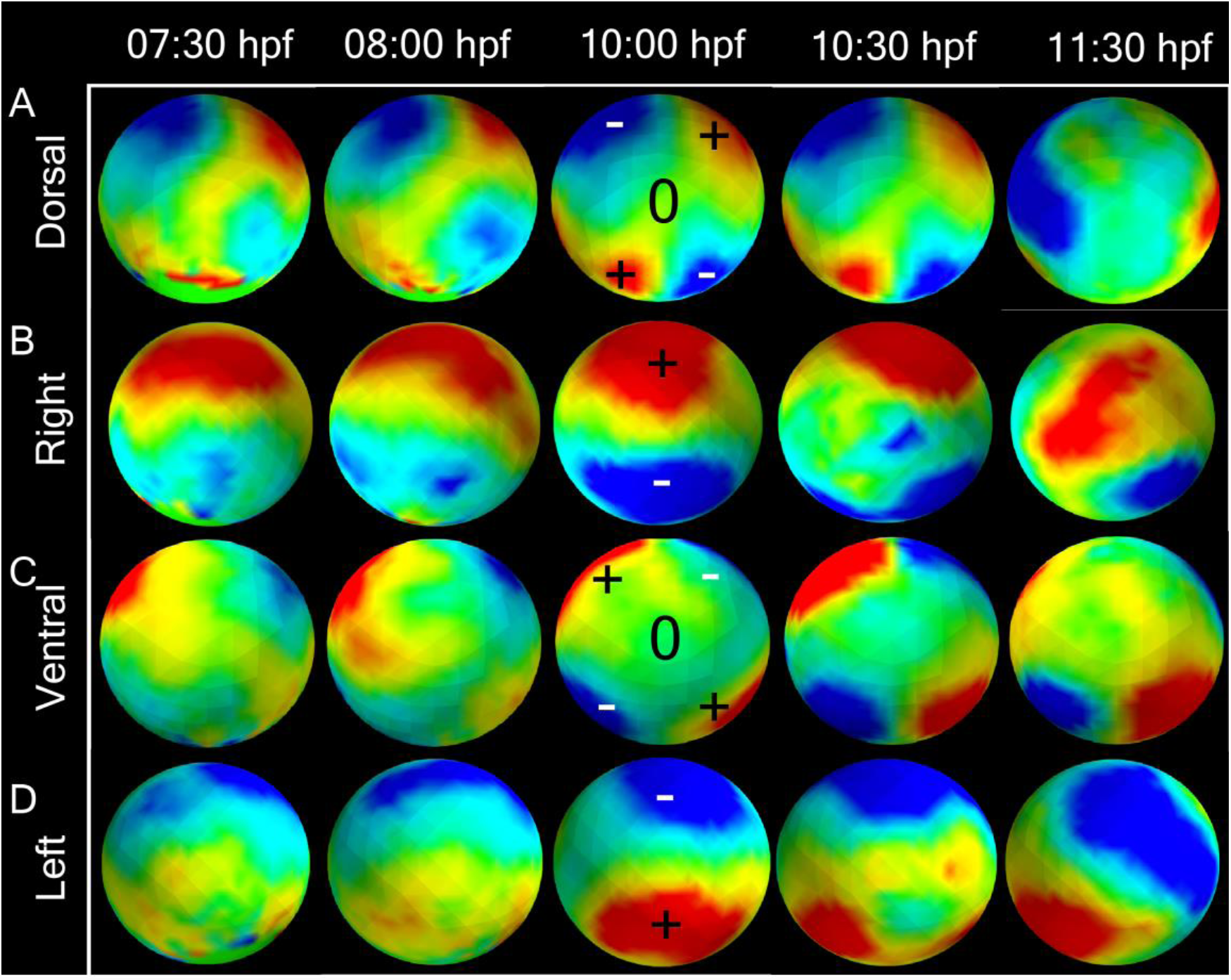
Positive (red) and negative (blue) rotational strain quantified by curl at 8 (75% epiboly), 9 (90% epiboly), 10 (bud, 100% epiboly), and 10.5 hpf (1-somite). In (A), (B), and (C), the embryo is oriented with anterior and posterior to the top and bottom. Positive curl is clockwise, negative curl is counter-clockwise. (A) Curl map of rotational strain over the dorsal view of the embryo. (B) Right side view of rotational strain of the embryo. (Dorsal is in the left side) (C) Curl map of rotational strain over the ventral view of the embryo. (D) Left side view of rotational strain of the embryo. (Dorsal is in the right side)

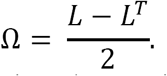

Positive or counter-clockwise rotation is red, negative or clockwise rotation is blue.

### Kymographs

The time evolution of strain is captured in a moving 20 min window from an annular region with a span of 0.38 radians. The axial, equatorial, or radial strain is averaged over domains along the equatorial circumference which bisects the dorsal (0°) and ventral (+/-180°) poles of the embryo or axial circumference in the AP/DV plane.

Author contributions
DB and PM designed the research, DB and JZ performed the research, STcontributed the GFP-histone construct, AK contributed the computational algorithm, DB, JZ, andPManalysed the data, DB, JZ, and PM wrote the paper, DB, JZ, AK, PM edited the paper

